# Poke And Delayed Drink Intertemporal Choice Task (POKE-ADDICT): an open-source behavioral apparatus for intertemporal choice testing in rodents

**DOI:** 10.1101/2023.09.08.556794

**Authors:** Andrea Stefano Moro, Daniele Saccenti, Alessia Seccia, A Mattia Ferro, Antonio Malgaroli, Jacopo Lamanna

**Affiliations:** Department of Psychology, Sigmund Freud University, Milan, Italy; Center for Behavioral Neuroscience and Communication (BNC), Vita-Salute San Raffaele University, Milan, Italy; Transcranial Magnetic Stimulation Unit, Italian Psychotherapy Clinics, Milan, Italy; Faculty of Psychology, Vita-Salute San Raffaele University, Milan, Italy; San Raffaele Turro, IRCCS Ospedale San Raffaele, Milan, Italy

## Abstract

Advancements in neuroscience research present opportunities and challenges, requiring substantial resources and funding. To address this, we describe here “Poke And Delayed Drink Intertemporal Choice Task (POKE-ADDICT)”, an open-source, versatile, and cost-effective apparatus for intertemporal choice testing in rodents. This allows quantification of delay discounting (DD), a cross-species phenomenon observed in decision making which provides valuable insights into higher-order cognitive functioning. In DD, the subjective value of a delayed reward is reduced as a function of the delay for its receipt. Using our apparatus, we implemented an effective intertemporal choice paradigm for the quantification of DD based on an adjusting delayed amount (ADA) algorithm using mango juice as a reward. Our paradigm requires limited training and can be directly translated to human subjects using monetary rewards. Our apparatus comprises a few 3D-printed parts and inexpensive electrical components, including a Raspberry Pi control unit. Furthermore, it is compatible with several *in vivo* procedures and the use of nose pokes instead of levers allows for faster task learning. Beside the main application described here, the apparatus can be further extended to implement other behavioral tests and protocols, including standard operant conditioning. In conclusion, we describe a versatile and cost-effective design based on Raspberry Pi that can support research in animal behavior, decision making and, more specifically, delay discounting.

## Introduction

Technological advancements in neuroscience research have brought forth both new opportunities and challenges. On the other hand, investigators require costly equipment and substantial funding to maintain competitiveness. Without adequate resources, the likelihood of conducting innovative studies becomes exceedingly low (Moro, 2022). Luckily, exploiting basic and established experimental approaches, such as Skinner boxes for studying animal behavior, can still produce valuable results. Nevertheless, also a very simple behavioral setup for rodents can represent a significant expense. In addition, commercial products often do not provide the required versatility for developing custom experimental protocols. To address these issues, we present an open-source, versatile, and cost-effective apparatus for conducting intertemporal choice testing in rodents. Among the various higher-order psychological functions, only a few are quantifiable and can be modeled in a mathematical manner. Delay discounting (DD) is a cross-species phenomenon observed in decision making where the subjective value of a delayed reward is reduced as a function of the delay for its receipt. DD is classically modelled using hyperbolic (Mazur, 1987) or exponential (Samuelson, 1937) functions which describe the tendency for more remote rewards to have less and less subjective value for their recipient (Odum et al., 2020). Indifference points, reflecting the subjective value of delayed outcomes (the amount of reward that, if given immediately, is considered equal to the delayed one) can be obtained through the intertemporal choice paradigm. Gauging the steepness of curves that mirror reward devaluation provides valuable information also in relation to clinical conditions, as DD is considered a transdiagnostic process (Amlung et al., 2019). Indeed, subjects affected by major depression (Pulcu et al., 2014), schizophrenia (Wang et al., 2018), borderline personality disorder (Krause-Utz et al., 2016), bipolar disorder (Urošević, Youngstrom, Collins, Jensen, & Luciana, 2016), bulimia nervosa (Kekic et al., 2016), and binge-eating disorder display steeper discounting curves compared to healthy controls (Manwaring, Green, Myerson, Strube, & Wilfley, 2011); whereas anorexic patients exhibit shallower reward devaluations (Lamanna et al., 2019; Steinglass et al., 2012).

DD also interacts with several other cognitive domains, including intelligence (Shamosh & Gray, 2008), working memory (Renda, Stein, & Madden, 2015) and episodic memory (Peters & Büchel, 2010). However, the causal relationships between psychopathology, cognitive dimensions, and DD remain unclear, highlighting the need for further research. Furthermore, the specific brain regions involved in this process are still not well understood (Moro, Saccenti, Ferro, et al., 2023). Dorsolateral and orbitofrontal portions of the prefrontal cortex, especially, configure among the best candidates for DD processing (He et al., 2016; Mar, Walker, Theobald, Eagle, & Robbins, 2011; Moro, Saccenti, Vergallito, et al., 2023; Nejati, Salehinejad, & Nitsche, 2018), but consistent causal evidences on such matter are still lacking. Considering these knowledge gaps and acknowledging the presence of DD in animals (Vanderveldt, Oliveira, & Green, 2016), we developed a custom-built apparatus for implementing intertemporal choice paradigms for rodents, potentially applicable for unveiling the neural mechanisms underlying DD. While existing literature presents various protocols for studying rodents (e.g. da Costa Araújo et al., 2009; Koot, Adriani, Saso, van den Bos, & Laviola, 2009; Mitchell, 2014; Richards, Mitchell, de Wit, & Seiden, 1997; Valencia-Torres et al., 2012; Wahab, Panlilio, & Solinas, 2018), to date, there is no standardized procedure for obtaining the discounting curve. Moreover, none of the existing protocols are suitable for laboratories with limited resources. To address this gap, our apparatus can be easily integrated into any operating chamber, was based on Raspberry Pi technology (which is widely available), three 3D-printed parts (two nose-pokes and one dish for liquid delivery) and a few inexpensive electrical components (see Materials section). Mango juice was selected as reward, given its pleasant and highly rewarding nature for rats, as it is calorically dense and rich in sugar (Lenoir, Serre, Cantin, & Ahmed, 2007). Additionally, we implemented an adjusting delayed amount two-choice task where the magnitude of the delayed reward is progressively modified, which allows for precise and reliable estimation of indifference points within a one-month timeframe. In this protocol, we opted to adjust the magnitude of the delayed reward, as it allows us to consider a broader quantitative range of responses without the constraint of a fixed larger reward. Importantly, procedures using either adjusting delay or adjusting reward algorithms produced no significant differences in terms of DD quantification (Holt, Green, & Myerson, 2012). Nevertheless, being open-source, the adopted algorithm can be modified as desired.

## Materials

### Apparatus

The behavioral experiments were conducted in a modular operant test chamber. Nevertheless, our components can be easily adapted to any kind of commercial or custom testing chamber provided minimal modifications. All 3D components were designed using Autodesk Fusion 360 (https://www.autodesk.com/products/fusion-360) and printed using a commercial low-cost 3D printer (MINGDA D2, www.3dmingdaofficial.com) and standard polylactide (PLA); design files are available at https://github.com/jacolamanna/POKE-ADDICT (GitHub). Electronic components are listed in Table 1. The total costs of materials (excluding the testing chamber) are below 50 €. Two 3D-printed nose-poke panels (Fig. 1a) were positioned in the opposite bottom corner of a wall, with the reward panel (Fig. 1b) placed in the center. The design of the nose-poke panels included an upper LED (blue or red) and a distance sensor (Mh-Sensor-Series, Flying-Fish) integrated into each hole (Table 1). The reward panel featured a dish for collecting mango juice and two tunnels/holes: one for liquid delivery and the other for disposal of excess juice and dish cleaning. These tunnels/holes were selectively connected to a semi-open bottle, allowing the mango juice to flow from its container into the dish via gravity, while the excess juice was directed towards a vacuum. The flow of juice was regulated by an electrovalve (Table 1), which enabled the administration of a specific number of drops during the trials. All electronic components, such as LEDs and distance sensors, were directly connected to the Raspberry Pi 3B+ pins, except for the electrovalve. The electrovalve, requiring a 12 V power supply, was connected to an external transformer and controlled through a Raspberry Pi pin via a Switch/Relay (Table 1). Figure 1c provides a summary of the apparatus.

**Table 1.** List of electronic components. The table displays the components usable for constructing the apparatus. The codes and products have been extracted from the website https://www.digikey.it/. As for the pinch solenoid valve, the one that, when closing, blocks the tube and the flow of mango juice, it is not available on the portal. However, it can be found on other websites for approximately € 30. While the distance sensor is available on many e-commerce websites, a package of 10 units can be found for approximately € 12.

| Item | Quantity | Price for unit (€) |
| --- | --- | --- |
| Raspberry Pi 3 Model B+ | 1 | 38.53 |
| LED (HL3P-BR60-J00DD) | 2 | 0.52 |
| Relay (G5LE-1 DC3) | 1 | 1.50 |
| Pinch Solenoid Valve ( SIRAI S104 01 EG W4 12V) | 1 | N.A. |
| Transformer (VEL05US120-US-JA) | 1 | 6.11 |
| Distance sensor (Mh-Sensor-Series, Flying-Fish) | 2 | N.A. |

**Figure 1.**
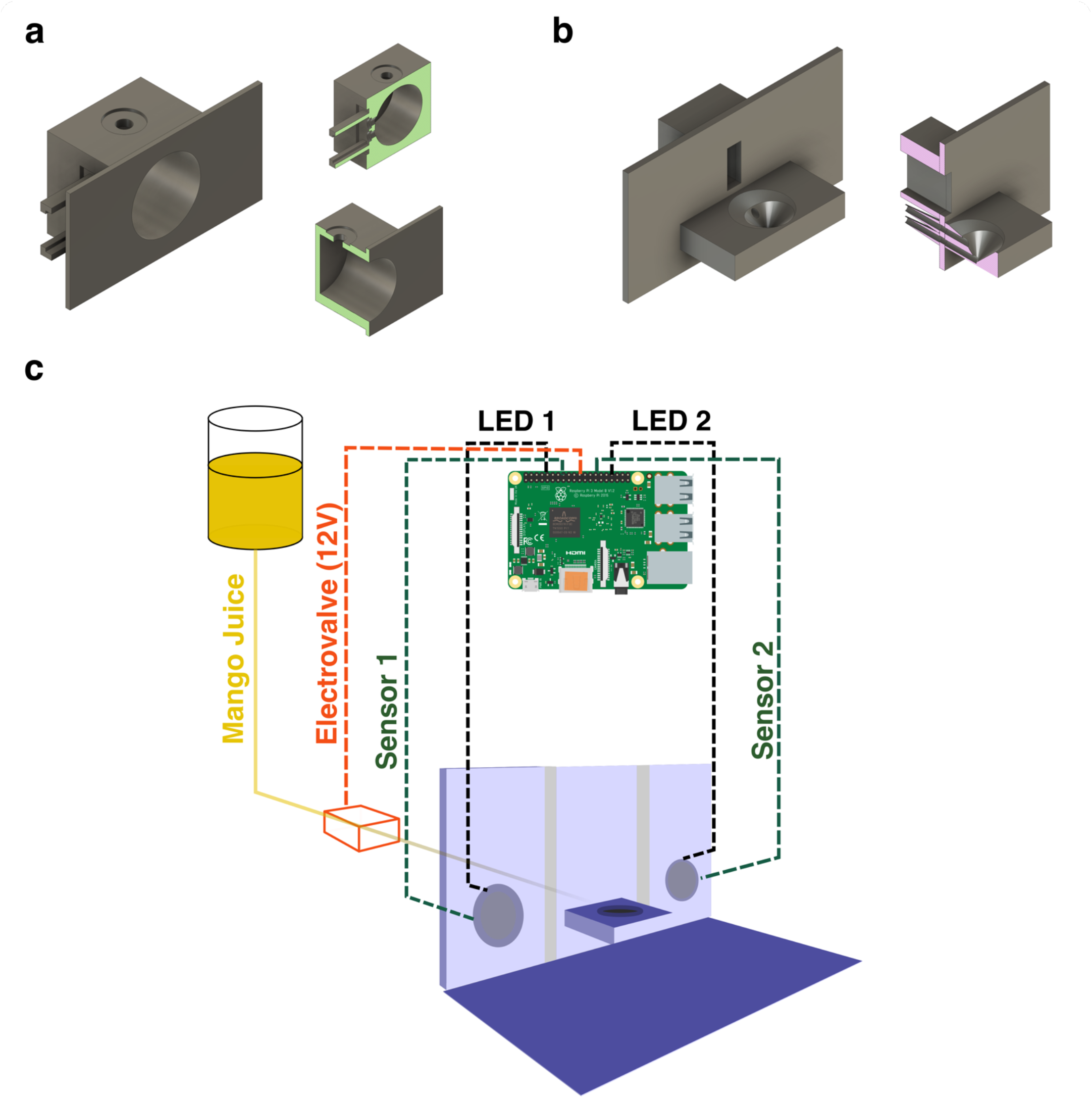
Schematic representation of the POKE-ADDICT apparatus. a) The rendering of the two Nose Poke ports (left). It is possible to observe the cavity to house the LED and the housing for the distance sensor. b) The rendering of the Reward Dish. c) The testing setup involves a Raspberry Pi 3b+ as the central component, which interfaces with the surrounding testing environment. The inputs to the system are received through two distance sensors (Mh-Sensor-Series, Flying-Fish) that are seamlessly integrated within the nose-poke holes. On the output side, the system employs a relay component to channel a 12 V power supply to the electrovalve. Additionally, two 3.3 LED indicators are positioned at the top of the nose-poke holes.

### Software

The experimental protocols were implemented using custom Python code that ran directly on the Raspberry Pi (all codes available at https://github.com/jacolamanna/POKE-ADDICT). Graphs and analyses were performed using R (https://www.r-project.org/).

### Subjects

The experiments were conducted on three Sprague-Dawley rats (175-200 g upon arrival; Charles River, Italy; 6 weeks of age). When not undergoing testing protocols, the rats were housed under a 12-hour light/12-hour dark cycle, with ad libitum access to food and water, and maintained at a constant temperature of 23 °C. All experiments were conducted during the light phase. All procedures were carried out in compliance with relevant guidelines and regulations. The protocols were approved by the Animal Care and Use Committee of San Raffaele Scientific Institute, in accordance with the guidelines set forth by the Italian Ministry of Health (IACUC 905). The study adheres to the ARRIVE guidelines for reporting.

### Protocol

Prior to conducting the “Adjusting Amount Reward Test,” the rats underwent training phases consisting of a “Habituation Phase” and a “Learning Phase.” The task was divided into four working days, corresponding to a week. The experimental manipulations were performed between 10 am and 6 pm as required. Before each session, food access was restricted to approximately 80% of the rats’ daily requirements.

#### Habituation Phase

During the first week, the animals were habituated to the experimenter and the testing cage. For the first two days, each rat was handled by the experimenter and placed inside the experimental cage for several minutes. Throughout the remainder of the week, the experimenter exposed the rodent to mango juice using a plastic syringe, delivering a few milliliters into the dish. An important objective of this phase was the habituation to the reward. For all the rats in our sample, this habituation occurred within the first week.

#### Learning Phase

During the second week, the rodents were trained on the following sequence: *i)* association between inserting the head into the nose poke and reward release, *ii)* recognition of the light in the nose poke as an indicator of willingness to perform the task, and *iii)* response to the flashing light as a cue to wait. These three points were introduced sequentially, respecting the order outlined. For the first point, the experimenter used an algorithm that activated both LEDs of the two nose pokes, and when the rat inserted its head, 10 drops of mango juice were suddenly released into the plate (Figure 2a). To facilitate the association between nose poke activation and reward release, a few drops of mango juice were manually placed inside the nose poke using a plastic syringe. The objective of the second point was achieved by modifying the algorithm, wherein the LED of one of the two nose pokes alternately lit up. The rat learned to insert its head into the illuminated nose poke to receive the reward of 10 drops of mango juice (Figure 2b).

**Figure 2.**
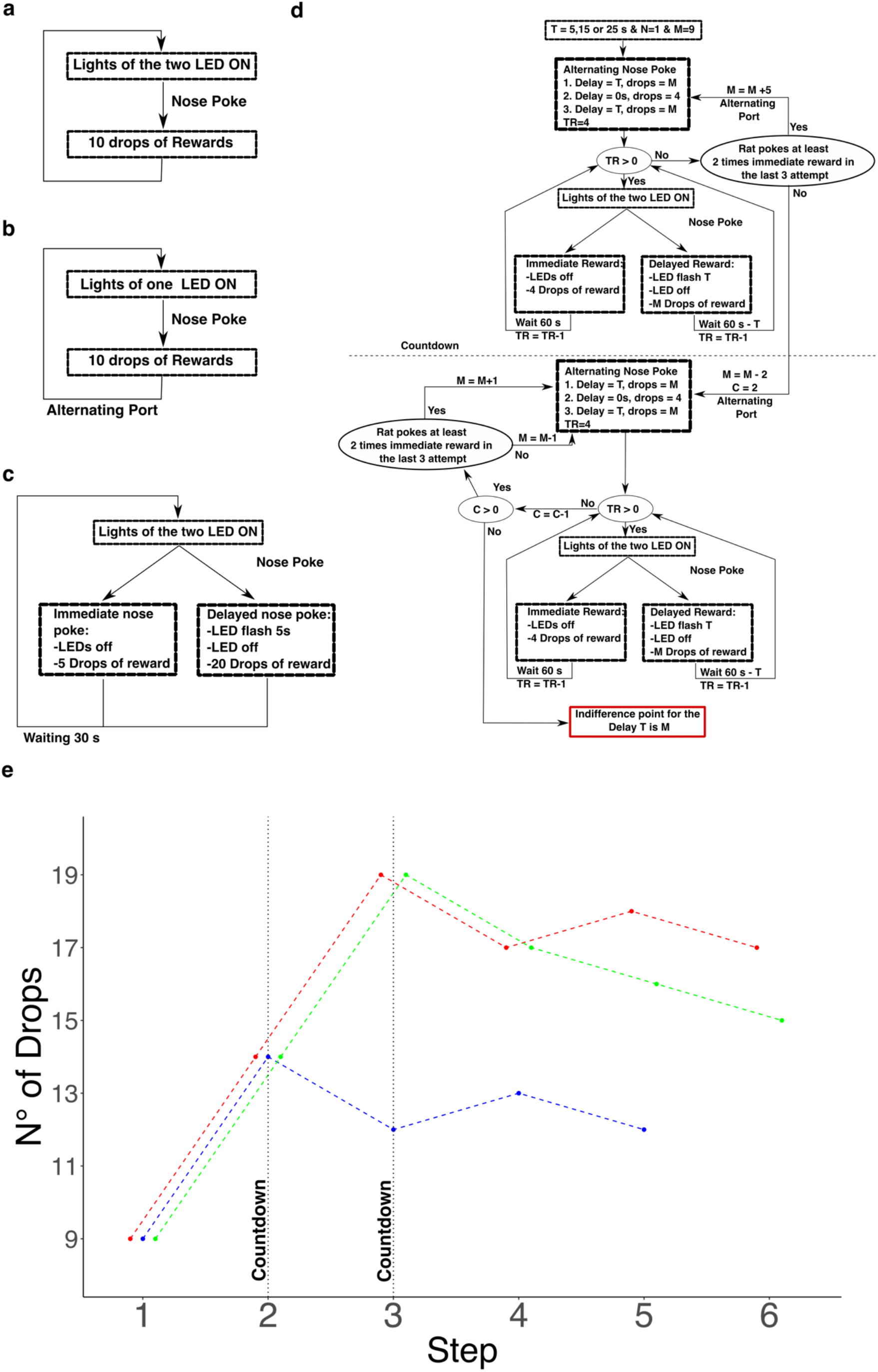
Training and testing algorithms. a-c) The flowcharts presented herein serve as schematic representations outlining the specific procedures employed during the Learning Phase of the experiment. d) Within the flowchart, a preconfigured scheme known as the Adjusting Amount Reward is delineated. This scheme governs the adjustment process for the reward magnitude. e) An illustrative example is provided within the algorithm, specifically highlighting a step where the delay is set to 15 seconds. The three distinct colors utilized in the representation correspond to the three individual rats involved in the study. On the y-axis, the computed magnitude of the reward is plotted according to the Adjusting Amount Reward algorithm, while the x-axis indicates the step associated with adjusting the quantity of drops. The vertical dashed line signifies the initiation of the countdown for convergence.

Regarding the last point, the training algorithm was further modified. One nose poke produced an immediate reward of 5 drops of mango juice, while the other nose poke led to a delayed reward of 20 drops after a 5-second delay. When the rat selected the nose poke with the delayed reward, the LED of the selected nose poke started flashing throughout the delay, while the LED of the other nose poke turned off. It is recommended to alternate the nose pokes, for example, after every 10 trials, for geting rid of any possible preference bias (Figure 2c).

Before proceeding to the next step, it is important to ensure that the rat performed each task within a short timeframe, i.e., less than 60 seconds.

#### Adjusting Delayed Amount Reward Test

From the third week onwards, we administered the testing protocols using the adjusting delayed amount two-choice task. Each rat underwent several blocks, where one nose poke was associated with an immediate reward (‘I’) of 4 drops, and the other nose poke was associated with a delayed but larger reward (‘D’) of 9 drops. Each block consisted of three forced-alternate trials (D, I, and D) followed by four free-choice trials. Before initiating the next trial, a 60-second inter-trial interval was initiated from the time of nose poking. At the end of each block, we considered the outcomes of the last three trials. If the rat consistently chose the immediate reward for all three trials, we added 5 drops to the delayed choice. If not, we initiated the “countdown” by subtracting 2 drops from the delayed choice. The nose poke positions were then inverted to prevent a preference bias.

For the subsequent block after the countdown had begun, we again considered the outcomes of the last three trials. If the rat chose the immediate reward at least twice, we added 1 drop to the delayed choice. Otherwise, we subtracted 1 drop for the future block. In the following block, the same procedure was applied, adding or subtracting 1 drop based on the outcomes of the last three trials. At this point, the indifference point was determined (Figure 2d). Once the animal reached the indifference point, we randomly changed the delay and repeated the process. We tested three different delays: 5, 15, and 25 seconds. An example of this process is illustrated in Figure 2e.

After testing a specific delay, the rat was typically replaced in the testing cage. Each delay was normally tested once per day.

Since the third week, when the animals were not yet expert in the task, the results were not considered reliable. Data collection started when the rat consistently approached the nose poke within a minute, which usually occurred during the fourth or fifth week (Figure 3).

**Figure 3.**
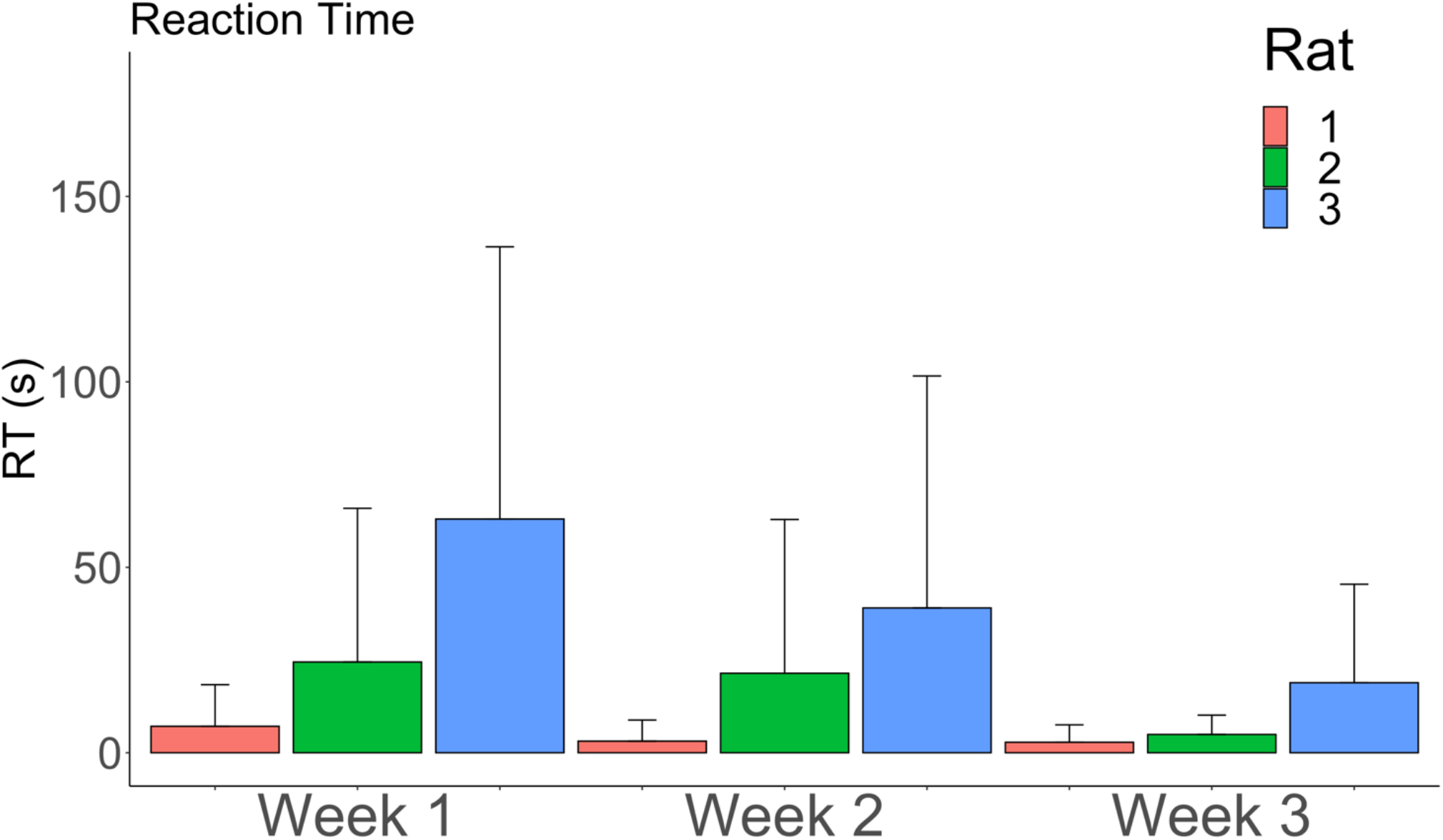
Reaction times. The response times exhibited by the rodents over the initial three-week period demonstrate a notable interindividual variability. However, it is observed that across all three rats, there is a consistent reduction in reaction times during the third week.

## Discussion

Investigating behavioral performance is a challenging task due to the difficulty in accurately measuring and quantifying the outcomes. In our manuscript, we presented a behavioral protocol to assess delay discounting, a psychological domain that can be quantitatively measured and modeled (Moro, Saccenti, Ferro, et al., 2023), and provided all details for the construction of an inexpensive, custom-built behavioral apparatus. The examination of intertemporal choice performance not only enhances our understanding of neuronal mechanisms and brain circuits involved but also foster the study of animal models of psychiatric disorders. Individuals with either substance or behavioral addiction, for instance, tend to choose smaller but sooner rewards (Albein-Urios, Martinez-González, Lozano, & Verdejo-Garcia, 2014; Amlung et al., 2019), whereas patients with anorexia nervosa exhibit a preference for delayed rewards (Steward et al., 2017). Greater discounting rates have been observed in rodents as well, e.g. in binge-eating rats when compared to controls, along with lisdexamfetamine reversing their reduced preference for larger later rewards (Vickers, Goddard, Brammer, Hutson, & Heal, 2017).

Importantly, our apparatus is compatible with various *in vivo* procedures, including electrophysiological, surgical, and pharmacological interventions. In particular, the administration of neuroactive chemicals able to block or enhance neuronal activity, e.g. tetrodotoxin or bicuculline, at specific brain sites, followed by the assessment of animals’ behavior, might contribute to understand the role of many untested regions in the mechanisms behind DD processes. By utilizing the modular operant test chamber and employing mango juice as the reward instead of solid food, we not only reduced costs and building complexity, but also achieved accurate and high-resolution quantification of the reward. This enhances the reliability when identifying indifference points. Additionally, the use of nose pokes instead of levers enables faster task learning, given the contingency of food reinforcement (Schindler, Thorndike, & Goldberg, 1993). Although several papers have employed adaptive algorithms (Mitchell, 2014; Wahab et al., 2018), the algorithm we propose exhibits high precision in determining the indifference points.

Another objective of our paper is to provide ideas for developing affordable and highly versatile experimental apparatus. The incorporation of electronic systems such as Raspberry Pi allows for a high degree of customization. Researchers can easily integrate new effectors and sensors, as well as create tailored algorithms for various behavioral protocols. Moreover, these systems are cost-effective, enabling low-budget laboratories to conduct competitive experiments.

A limitation of our study is the small number of animals used for testing the apparatus. However, similar works have also employed a limited number of rodents (Koot et al., 2009), and increasing the sample size would not sensibly contribute to testifying the functionality and advantages of our apparatus and protocol. Importantly, by observing the behavior of tested rats (Figure 4), we confirmed the well-established finding that increasing reward delays decrease the subjective value of the reward (Myerson, Green, & Warusawitharana, 2001).

**Figure 4.**
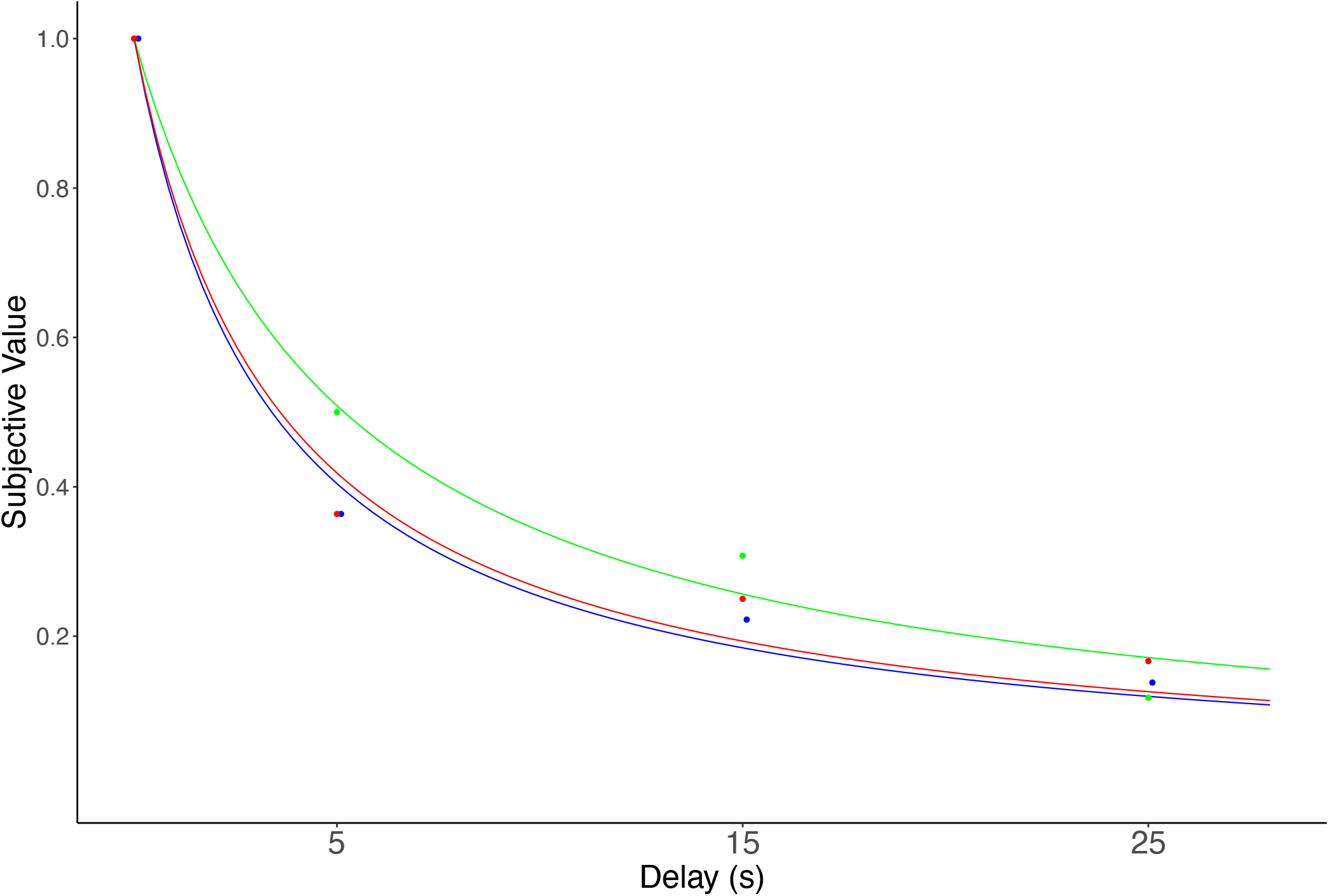
Indifference points and discounting curves. The plot illustrates estimated indifferent points obtained using POKE-ADDICT within a single experimental session and fitted discounting curves. As the delay increases, estimated subjective value reduces, as expected. Each distinct color within the graph represents the performance of an individual rat. We fitted the subjective value points with a hyperbolic function (Subjective Value = 1 / (1 + k * Delay)). Indeed, the data reflects the typical hyperbolic pattern found for delay discounting.

## References

Albein-Urios, N., Martinez-González, J. M., Lozano, Ó., & Verdejo-Garcia, A. (2014). Monetary delay discounting in gambling and cocaine dependence with personality comorbidities. Addictive Behaviors, 39(11), 1658–1662. 10.1016/j.addbeh.2014.06.001

Amlung, M., Marsden, E., Holshausen, K., Morris, V., Patel, H., Vedelago, L., … McCabe, R. E. (2019). Delay Discounting as a Transdiagnostic Process in Psychiatric Disorders: A Metaanalysis. JAMA Psychiatry, 76(11), 1176–1186. 10.1001/jamapsychiatry.2019.2102

da Costa Araújo, S., Body, S., Hampson, C. L., Langley, R. W., Deakin, J. F. W., Anderson, I. M.,… Szabadi, E. (2009). Effects of lesions of the nucleus accumbens core on inter-temporal choice: Further observations with an adjusting-delay procedure. Behavioural Brain Research, 202(2), 272–277. 10.1016/j.bbr.2009.04.003

He, Q., Chen, M., Chen, C., Xue, G., Feng, T., & Bechara, A. (2016). Anodal stimulation of the left DLPFC increases IGT scores and decreases delay discounting rate in healthy males. Frontiers in Psychology, 7(SEP), 1–9. 10.3389/fpsyg.2016.01421

Holt, D. D., Green, L., & Myerson, J. (2012). Estimating the subjective value of future rewards: Comparison of adjusting-amount and adjusting-delay procedures. Behavioural Processes, 90(3), 302–310. 10.1016/j.beproc.2012.03.003

Kekic, M., Bartholdy, S., Cheng, J., McClelland, J., Boysen, E., Musiat, P., … Schmidt, U. (2016). Increased temporal discounting in bulimia nervosa. International Journal of Eating Disorders, 49(12), 1077–1081. 10.1002/eat.22571

Koot, S., Adriani, W., Saso, L., van den Bos, R., & Laviola, G. (2009). Home cage testing of delay discounting in rats. Behavior Research Methods, 41(4), 1169–1176. 10.3758/BRM.41.4.1169

Krause-Utz, A., Cackowski, S., Daffner, S., Sobanski, E., Plichta, M. M., Bohus, M., … Schmahl, C. (2016). Delay discounting and response disinhibition under acute experimental stress in women with borderline personality disorder and adult attention deficit hyperactivity disorder. Psychological Medicine, 46(15), 3137–3149. 10.1017/S0033291716001677

Lamanna, J., Sulpizio, S., Ferro, M., Martoni, R., Abutalebi, J., & Malgaroli, A. (2019). Behavioral assessment of activity-based-anorexia: how cognition can become the drive wheel. Physiology & Behavior, 202, 1–7. 10.1016/j.physbeh.2019.01.016

Lenoir, M., Serre, F., Cantin, L., & Ahmed, S. H. (2007). Intense Sweetness Surpasses Cocaine Reward. PLoS ONE, 2(8), e698. 10.1371/journal.pone.0000698

Manwaring, J. L., Green, L., Myerson, J., Strube, M. J., & Wilfley, D. E. (2011). Discounting of Various Types of Rewards by Women with and Without Binge Eating Disorder: Evidence for General Rather Than Specific Differences. The Psychological Record, 61(4), 561–582. 10.1007/BF03395777

Mar, A. C., Walker, A. L. J., Theobald, D. E., Eagle, D. M., & Robbins, T. W. (2011). Dissociable Effects of Lesions to Orbitofrontal Cortex Subregions on Impulsive Choice in the Rat. The Journal of Neuroscience, 31(17), 6398–6404. 10.1523/JNEUROSCI.6620-10.2011

Mazur, J. E. (1987). An adjusting procedure for studying delayed reinforcement. In Quantitative Analyses of Behavior, Vol. 5. The effect of delay and of intervening events on reinforcement value. (pp. 55–73). Hillsdale, NJ, US: Lawrence Erlbaum Associates, Inc.

Mitchell, S. H. (2014). Assessing delay discounting in mice. Current Protocols in Neuroscience, (SUPPL.66), 1–12. 10.1002/0471142301.ns0830s66

Moro, A. S. (2022). A Concrete Recipe to Reinvent and Innovate the Bachelor ’ s Program : Free Choice of Courses and Hackathon -Based Teaching. Human Arenas, (0123456789). 10.1007/s42087-022-00301-x

Moro, A. S., Saccenti, D., Ferro, M., Scaini, S., Malgaroli, A., & Lamanna, J. (2023). Neural Correlates of Delay Discounting in the Light of Brain Imaging and Non-Invasive Brain Stimulation : What We Know and What Is Missed. Brain Science.

Moro, A. S., Saccenti, D., Vergallito, A., Scaini, S., Malgaroli, A., Ferro, M., & Lamanna, J. (2023). Transcranial direct current stimulation (tDCS) over the orbitofrontal cortex reduces delay discounting. (August), 1–11. 10.3389/fnbeh.2023.1239463

Myerson, J., Green, L., & Warusawitharana, M. (2001). AREA UNDER THE CURVE AS A MEASURE OF DISCOUNTING. Journal of the Experimental Analysis of Behavior, 76(2), 235–243. 10.1901/jeab.2001.76-235

Nejati, V., Salehinejad, M. A., & Nitsche, M. A. (2018). Interaction of the Left Dorsolateral Prefrontal Cortex (l-DLPFC) and Right Orbitofrontal Cortex (OFC) in Hot and Cold Executive Functions: Evidence from Transcranial Direct Current Stimulation (tDCS). Neuroscience, 369, 109–123. 10.1016/j.neuroscience.2017.10.042

Odum, A. L., Becker, R. J., Haynes, J. M., Galizio, A., Frye, C. C. J., Downey, H., … Perez, D. M. (2020). Delay discounting of different outcomes: Review and theory. Journal of the Experimental Analysis of Behavior, 113(3), 657–679. 10.1002/jeab.589

Peters, J., & Büchel, C. (2010). Episodic Future Thinking Reduces Reward Delay Discounting through an Enhancement of Prefrontal-Mediotemporal Interactions. Neuron, 66(1), 138–148. 10.1016/j.neuron.2010.03.026

Pulcu, E., Trotter, P. D., Thomas, E. J., McFarquhar, M., Juhasz, G., Sahakian, B. J., … Elliott, R. (2014). Temporal discounting in major depressive disorder. Psychological Medicine, 44(9), 1825–1834. 10.1017/S0033291713002584

Renda, C. R., Stein, J. S., & Madden, G. J. (2015). Working-memory training: Effects on delay discounting in male long evans rats. Journal of the Experimental Analysis of Behavior, 103(1), 50–61. 10.1002/jeab.115

Richards, J. B., Mitchell, S. H., de Wit, H., & Seiden, L. S. (1997). DETERMINATION OF DISCOUNT FUNCTIONS IN RATS WITH AN ADJUSTING-AMOUNT PROCEDURE. Journal of the Experimental Analysis of Behavior, 67(3), 353–366. 10.1901/jeab.1997.67-353

Samuelson, P. A. (1937). A Note on Measurement of Utility. The Review of Economic Studies, 4(2), 155. 10.2307/2967612

Schindler, C. W., Thorndike, E. B., & Goldberg, S. R. (1993). Acquisition of a nose-poke response in rats as an operant. Bulletin of the Psychonomic Society, 31(4), 291–294. 10.3758/BF03334932

Shamosh, N. A., & Gray, J. R. (2008). Delay discounting and intelligence: A meta-analysis. Intelligence, 36(4), 289–305. 10.1016/j.intell.2007.09.004

Steinglass, J. E., Figner, B., Berkowitz, S., Simpson, H. B., Weber, E. U., & Walsh, B. T. (2012). Increased capacity to delay reward in anorexia nervosa. Journal of the International Neuropsychological Society, 18(4), 773–780. 10.1017/S1355617712000446

Steward, T., Mestre-Bach, G., Vintró-Alcaraz, C., Agüera, Z., Jiménez-Murcia, S., Granero, R., & Fernández-Aranda, F. (2017). Delay Discounting of Reward and Impulsivity in Eating Disorders: From Anorexia Nervosa to Binge Eating Disorder. European Eating Disorders Review, 25(6), 601–606. 10.1002/erv.2543

Urošević, S., Youngstrom, E. A., Collins, P., Jensen, J. B., & Luciana, M. (2016). Associations of age with reward delay discounting and response inhibition in adolescents with bipolar disorders. Journal of Affective Disorders, 190, 649–656. 10.1016/j.jad.2015.11.005

Valencia-Torres, L., Olarte-Sánchez, C. M., da Costa Araújo, S., Body, S., Bradshaw, C. M., & Szabadi, E. (2012). Nucleus accumbens and delay discounting in rats: evidence from a new quantitative protocol for analysing inter-temporal choice. Psychopharmacology, 219(2), 271– 283. 10.1007/s00213-011-2459-1

Vanderveldt, A., Oliveira, L., & Green, L. (2016). Delay discounting: Pigeon, rat, human—does it matter? Journal of Experimental Psychology: Animal Learning and Cognition, 42(2), 141– 162. 10.1037/xan0000097

Vickers, S. P., Goddard, S., Brammer, R. J., Hutson, P. H., & Heal, D. J. (2017). Investigation of impulsivity in binge-eating rats in a delay-discounting task and its prevention by the damphetamine prodrug, lisdexamfetamine. Journal of Psychopharmacology, 31(6), 784–797. 10.1177/0269881117691672

Wahab, M., Panlilio, L. V, & Solinas, M. (2018). A modified self-adjusting delay discounting procedure for the study of choice impulsivity in rats A modified self-adjusting delay discounting proce-dure for the study of choice impulsivity in rats. Psychopharmacology, 235(7). 10.1007/s00213-018-4911-yï

Wang, L., Jin, S., He, K., Chen, X., Ji, G., Bai, X., … Wang, K. (2018). Increased delayed reward during intertemporal decision-making in schizophrenic patients and their unaffected siblings. Psychiatry Research, 262, 246–253. 10.1016/j.psychres.2017.12.040

